# Lifelong musical activity is associated with multi-domain cognitive and brain benefits in older adults

**DOI:** 10.1101/2021.09.15.460202

**Authors:** Adriana Böttcher, Alexis Zarucha, Theresa Köbe, Malo Gaubert, Angela Höppner, Klaus Fabel, Slawek Altenstein, Claudia Bartels, Katharina Buerger, Peter Dechent, Laura Dobisch, Michael Ewers, Klaus Fliessbach, Silka Dawn Freiesleben, Ingo Frommann, John Dylan Haynes, Daniel Janowitz, Ingo Kilimann, Luca Kleineidam, Christoph Laske, Franziska Maier, Coraline Metzger, Matthias Munk, Robert Perneczky, Oliver Peters, Josef Priller, Boris-Stephan Rauchmann, Nina Roy, Klaus Scheffler, Anja Schneider, Annika Spottke, Stefan J. Teipel, Jens Wiltfang, Steffen Wolfsgruber, Renat Yakupov, Emrah Duzel, Frank Jessen, Sandra Röske, Michael Wagner, Gerd Kempermann, Miranka Wirth, for the DELCODE study group

**Affiliations:** German Center for Neurodegenerative Diseases (DZNE), Dresden, Germany; Cognitive Neurophysiology, Department of Child and Adolescent Psychiatry, Faculty of Medicine, Technische Universität Dresden, Dresden, Germany; CRTD – Center for Regenerative Therapies, Technische Universität Dresden, Dresden, Germany; German Center for Neurodegenerative Diseases (DZNE), Berlin, Germany; Department of Psychiatry, Campus Benjamin Franklin, Charité – Universitätsmedizin Berlin, Berlin, Germany; Department of Psychiatry and Psychotherapy, University Medical Center, University of Goettingen, Goettingen, Germany; Institute for Stroke and Dementia Research (ISD), University Hospital, LMU Munich, Munich, Germany; German Center for Neurodegenerative Diseases (DZNE), Munich, Germany; MR-Research in Neurology and Psychiatry, Georg-August-University Göttingen, Goettingen, Germany; German Center for Neurodegenerative Diseases (DZNE), Magdeburg, Germany; Institute of Cognitive Neurology and Dementia Research (IKND), Otto-von-Guericke University, Magdeburg, Germany; Institute for Clinical Radiology, LMU Munich, Munich, Germany; German Center for Neurodegenerative Diseases (DZNE), Bonn, Germany; Department for Neurodegenerative Diseases and Geriatric Psychiatry, University Hospital Bonn, Bonn, Germany; Bernstein Center for Computational Neuroscience, Charité — Universitätsmedizin, Berlin, Germany; German Center for Neurodegenerative Diseases (DZNE), Rostock, Germany; Department of Psychosomatic Medicine, Rostock University Medical Center, Rostock; German Center for Neurodegenerative Diseases (DZNE), Tuebingen, Germany; Section for Dementia Research, Hertie Institute for Clinical Brain Research and Department of Psychiatry and Psychotherapy, University of Tuebingen, Tuebingen, Germany; Department of Psychiatry, University of Cologne, Medical Faculty, Cologne, Germany; Department of Psychiatry and Psychotherapy, Otto-von-Guericke University, Magdeburg, Germany; Munich Cluster for Systems Neurology (SyNergy) Munich, Munich, Germany; Ageing Epidemiology Research Unit (AGE), School of Public Health, Imperial College London, London, UK; Department of Psychiatry and Psychotherapy, University Hospital, LMU Munich, Munich, Germany; Department of Psychiatry and Psychotherapy, Klinikum rechts der Isar, Technical University Munich, Munich, Germany; Department for Biomedical Magnetic Resonance, University of Tuebingen, Tuebingen, Germany; Department of Neurology, University Hospital Bonn, Bonn, Germany; German Center for Neurodegenerative Diseases (DZNE), Goettingen, Germany; Neurosciences and Signaling Group, Institute of Biomedicine (iBiMED), Department of Medical Sciences, University of Aveiro, Aveiro, Portugal; Excellence Cluster on Cellular Stress Responses in Aging-Associated Diseases (CECAD), University of Cologne, Cologne, Germany

**Keywords:** cognitive reserve, resilience, prevention, brain plasticity, instrument playing

## Abstract

Regular musical activity as a highly-stimulating lifestyle activity is proposed to be protective against age-related cognitive decline and Alzheimer’s disease (AD). This study investigated associations between lifelong regular musical instrument playing, late-life cognitive abilities and brain morphology in older adults. We show that musical activity over the life course is associated with better global cognition, working memory, executive functions, language, and visuospatial abilities accounting for reserve proxies. Playing music is not significantly associated with gray matter volume in regions most affected by aging and AD. Selectively in the musically active participants, multi-domain cognitive abilities were enhanced with preserved gray matter volume in frontal and temporal regions. Our correlational findings suggest that playing a musical instrument may improve the recruitment of existing brain resources to facilitate late-life cognitive capacities. We propose that engaging in regular musical activity could serve as a low-threshold multimodal enrichment strategy that may promote cognitive resilience in advanced age.

## 2 Introduction

Healthy lifestyle activities are proposed to enhance brain and cognitive resilience in older adults ^1^ through multiple neuroprotective pathways ^2-4^ and may thereby offer protection against age-related neurodegenerative diseases, such as Alzheimer’s disease (AD). Among others, regular musical activity, such as playing an instrument, has been associated with reduced risk of developing dementia ^5,6^. To advance targeted intervention strategies, it is important to delineate cognitive benefits and underlying brain mechanisms associated with musical activity in advanced age.

Musical activity is suggested to share communalities with the concept of environmental enrichment ^7,8^, shown to promote far-reaching neurobiological and behavioral benefits in animal models ^9^. Playing a musical instrument entails complex skills involving the simultaneous perception and integration of motor, sensory, cognitive, emotional, and social stimulations, thought to facilitate beneficial brain plasticity ^10^. Consistently, there is evidence indicating that playing music could preserve higher-order cognitive abilities in older adults ^11,12^. In this population, benefits of regular musical activity have been shown to transfer to multiple cognitive domains that typically decline with higher age, including executive functions, attention, language, visuospatial as well as memory abilities ^13-17^. Together these findings imply that complex multimodal stimulation, as inherent to playing music, might help retain cognitive capacities in late life.

Comparatively little is, however, known about the neurobiological underpinnings of regular musical activity in older adults ^18^. Studies that have investigated brain correlates of playing music in young and middle-aged cohorts suggest that this activity leads to plasticity, as reflected in measurable volume increases in distributed brain areas ^19,20^. Those areas comprise multisensory frontal, partial, and temporal regions ^21-23^, which are strongly affected by healthy and pathological aging ^24-26^. Regularly participating in musical activity may also be protective for the hippocampus. In young musicians compared to controls, musical activity is associated with enhanced volume and functions of the hippocampus ^27-29^. In older adults, there appears to be a volume increase in frontal and temporal areas associated with playing music ^30^. Such benefits in brain resources may contribute to better late-life cognitive abilities associated with this lifestyle activity.

Overall, the existing findings propose that regular musical activity may protect brain and cognitive health via multiple pathways. There might be a boost in functional brain capacities and/or an increase in structural brain resources, both of which may help counteract neuropathological burden in advanced age ^31^. To shed light onto these mechanisms, this cross-sectional study investigated potential cognitive and brain benefits of playing music during the entire life in the older population. Taking into account reserve proxies of educational attainment, crystallized intelligence, socioeconomic status (SES), and physical activity, we hypothesized that lifelong regular musical activity is associated with better late-life cognitive abilities in multiple domains as well as larger gray matter volume (GMV), particularly in regions affected by healthy and pathological aging. We further investigated, whether playing an instrument has a positive influence on the association between regional brain structure and cognitive performance in older adults, proposed to convey cognitive resilience in advanced age.

## 3 Material and methods

### 3.1 Overall design of the DELCODE study

The data used in this study were obtained from the DZNE-Longitudinal Cognitive Impairment and Dementia cohort. The detailed study protocol can be found in a previous report ^32^. In brief, the DELCODE cohort was set up to recruit 1000 participants at baseline with five groups of participants. Specifically, these groups are healthy controls (HC), first-degree relatives of AD patients (family history, FH) as well as participants with subjective cognitive decline (SCD), mild cognitive impairment (MCI), and mild AD dementia. At baseline assessment, all participants received extensive clinical, neuropsychological, and behavioral assessments. To minimize site-effects and ensure high data quality, assessment protocols were standardized across sites using Standard Operating Procedures (SOP). Post-scanning MRI image quality assessments were conducted by the DZNE Magdeburg. The DELCODE study protocol agreed with ethical principles for human experimentation in accordance with the Declaration of Helsinki. At each participating study sites, the protocol was approved by the local ethical committees. All participants gave their written informed consent. DELCODE was registered at the German Clinical Trials Register (DRKS00007966; April 5, 2015).

### 3.2 Participants

In the present study, cognitively healthy participants (HC, FH, and SCD) were included and merged across the three groups to increase the final sample size. Recruitment procedures including inclusion and exclusion criteria are described in detail elsewhere ^32^. In brief, all participants were aged ≥ 60 years, German speaking, able to provide informed consent and had a study partner serving as an informant. Normal cognitive function was defined as a test performance within -1.5 standard deviations of age-, sex- and education-adjusted norms on all subtests of the Consortium to Establish a Registry of Alzheimer’s Disease (CERAD) test battery ^49^. Exclusion criteria for HC, FH, and SCD were comprised of medical conditions including current or past major medical, neurological, or psychiatric disorders. Presence of SCD was defined by subjectively reported decline in cognitive functioning with concerns ^50^. Diagnostic criteria for MCI and mild AD dementia are provided in Jessen et al. (2018).

The DELCODE baseline dataset (total: *n* = 1079) was used to select a subset of participants into the present study as follows (see Figure 1): At the time of our analysis, data from 943 participants with a structural cranial magnetic resonance imaging (MRI) assessment at baseline were available. Of these participants, cognitively healthy participants were selected (i.e., HC, SCD, FH, total: *n* = 678). Afterwards, participants who reported regular musical activity across the life span (i.e., group of interest) and participants with no musical activity during life (i.e., control group) were identified (total: *n* = 429; for methodological details see below). Finally, we matched the control group and included only participants with complete datasets regarding variables of interest, resulting in a final sample of *n* = 140 (for methodological details see below).

**Figure 1:**
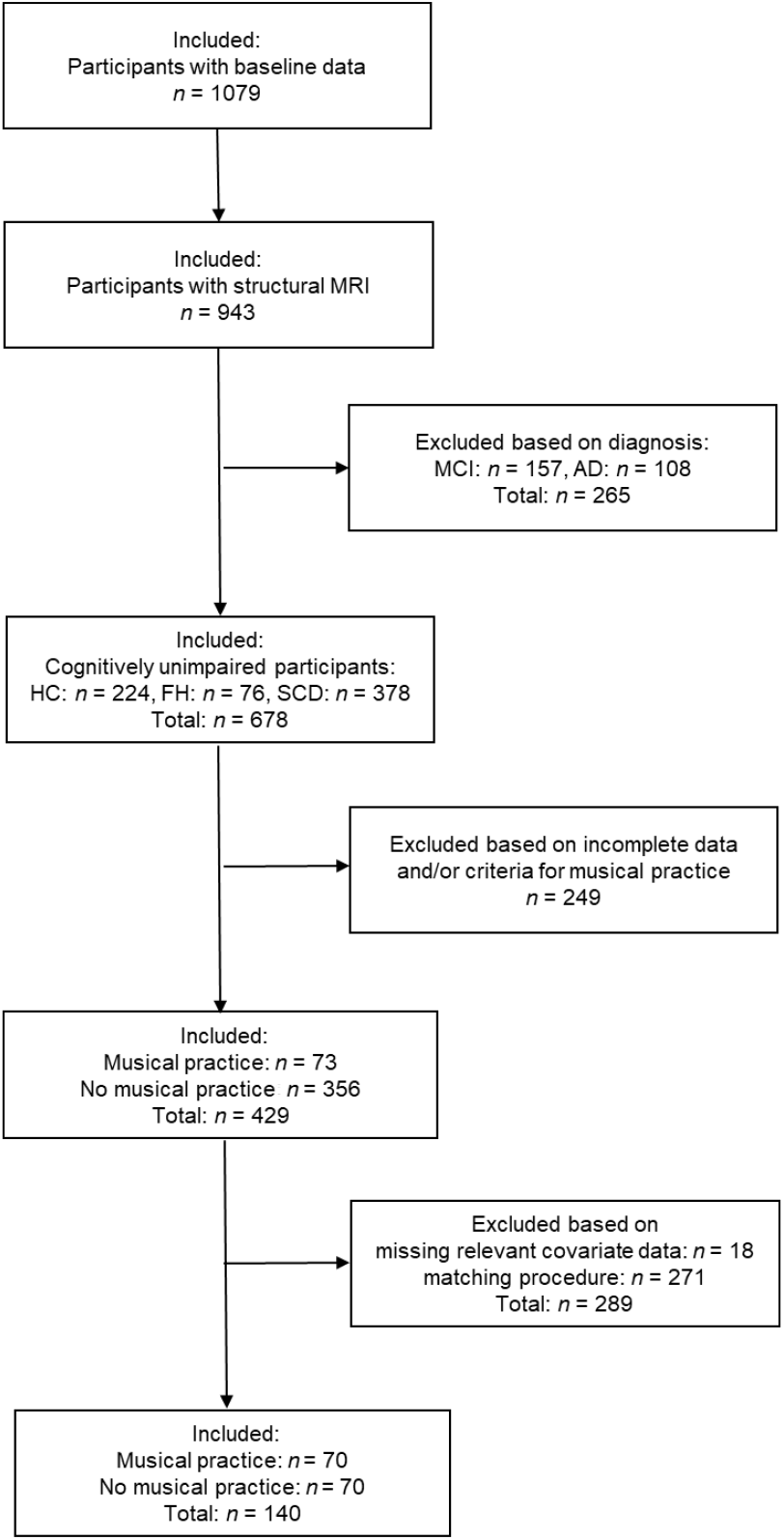
Participant selection flowchart. The graph displays the selection procedure from the DELCODE database. ***Key:*** HC, healthy controls; FH, family history of AD; AD, Alzheimer’s disease; MCI, mild cognitive impairment; Magnetic resonance imaging (MRI).

**Figure 2:**
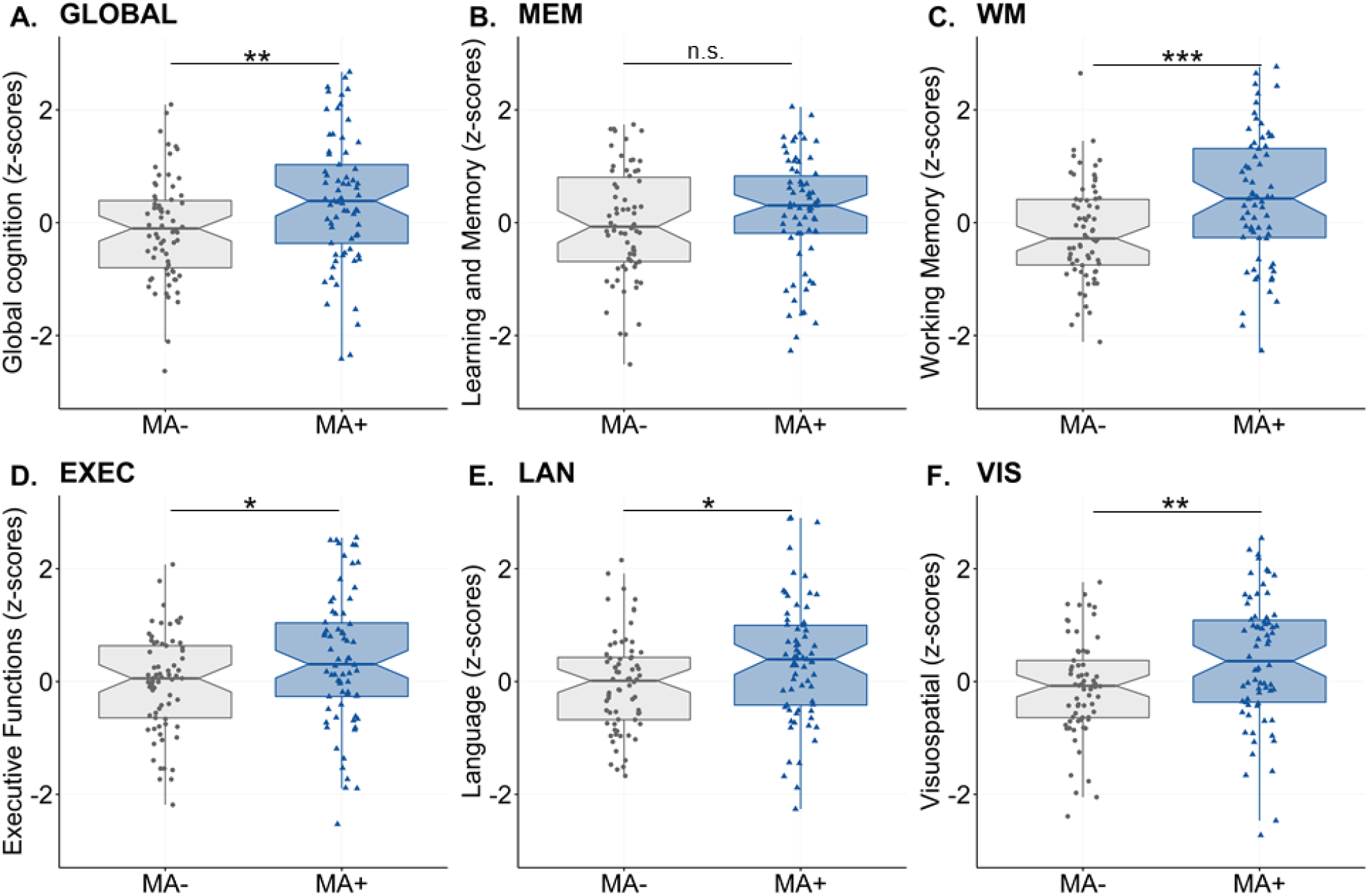
Main effect of lifelong musical activity on cognitive abilities. Significant group differences were found for global cognition (A, GLOBAL, working memory (C, WM), executive function (D, EXEC), language (E, LAN) and visuospatial abilities (F, VIS). These multi-domain cognitive abilities were enhanced for participants with lifelong musical activity (MA+, blue) compared to controls (no musical activity across lifespan, MA-, gray). The association was not significant for the learning and memory composite (B, MEM). Boxplots display unadjusted data with individual data points. The “notch” shows the median with 95% confidence intervals and interquartile range with lower (25th) and upper percentiles (75th). Significance levels (uncorrected): \*\*\**p* < 0.001, \*\**p* < 0.01, \**p* < 0.05. ***Key:*** MA+, musical activity; MA-, no musical activity.

### 3.3 Measurements

#### 3.3.1 Measurement of musical activity

Musical activity across lifespan was assessed using the Lifetime of Experiences Questionnaire ^47^, adapted for the German population ^51^. Details on the LEQ and the coding scheme used to assess lifelong regular musical activity are provided in the supplementary material. In brief, the self-reported questionnaire measures educational, occupational, and leisure activities across three life periods (young adulthood: 13 – 30 years, mid-life: 30 – 65 years, and late-life: 65 years onwards). One self-reported lifestyle activity inquired by the LEQ was the frequency of playing a musical instrument and this information was used to operationalize musical activity across the lifespan. Similar to a previous study ^33^, we constructed a variable that was comprised of two groups: (1) The musical activity group (group of interest) included those participants that were musically active in all life periods and reported regular musical activity (2 times per month or more) in at least one life period. (2) The no musical activity or control group included participants that reported to never have played a musical instrument in any of the life periods.

#### 3.3.2 Measures of cognitive abilities

Cognitive functioning was assessed using latent factors over five cognitive domains, namely (1) learning and memory, (2) working memory, (3) executive functions and mental processing speed, (4) language, and (5) visuospatial abilities, created by summarizing cognitive tests from the extensive neuropsychological test battery in the overall DELCODE cohort as described previously ^52^. In brief, Wolfsgruber et al. (2020) used confirmatory factor analyses (CFA) to extract the factor structure using data from the extensive neuropsychological test battery applied during baseline assessment. Additionally, a global cognitive performance score was calculated by taking the mean of the five cognitive scores ^52^. For the present analysis, we used performance measures for global cognition and the five cognitive domain scores. Each cognitive score was z-transformed using the selected DELCODE neuroimaging sample including HC, FH, and SCD participants.

#### 3.3.3 MRI acquisition and processing

MRI data were acquired using Siemens MRI scanners (Siemens, Erlangen, Germany), including three TIM Trio systems, four Verio systems, one Skyra system, and one Prisma system. The extensive MRI protocol of the DELCODE study is described elsewhere ^32^. For the present analysis, we used T1-weighted images (i.e., 3D GRAPPA PAT 2, 1 mm^3^ isotropic, 256 × 256 px, 192 slices, sagittal, ∼ 5min, TR 2500 ms, TE 4.33 ms, TI 110 ms, FA 7°) and T2-weighted images (i.e., 0.5 × 0.5 × 1.5 mm^3^, 384 × 384 px, 64 slices, orthogonal to hippocampal long axis, ∼12 min, TR 3500 ms, TE 353 ms, optimized for volumetric assessment of the medial temporal lobe). All scans underwent quality assessment provided by the DZNE imaging network (iNET, Magdeburg).

Regional GMV analysis was conducted in pre-selected regions-of-interest (ROI), robustly affected by healthy and pathological aging due to AD ^24-26^. Based on these findings, we chose two regions, that is, the frontal lobe and the hippocampus. For each of these ROIs, we used regional volume measures provided in the DELCODE database, as described previously ^53^. In brief, structural MRI images were segmented in native space using an automated cortical parcellation pipeline ^54^ implemented in FreeSurfer (version 6.0, http://surfer.nmr.mgh.harvard.edu/) and an advanced segmentation tool ^55^ to derive ROI-based GMV. To enhance reliability, image segmentation was based on T1-weighted and high-resolution T2-weighted images. Left and right hippocampal volume were summed for a measure of the overall hippocampal volume. Frontal volume was calculated as the sum over left and right frontal ROI of the Desikan atlas following the procedure described elsewhere ^56^. In addition, cortical GMV was evaluated as a global measure of brain integrity. Regional GMV measures were adjusted for total intracranial volume (TIV), as estimated using FreeSurfer ^57^, using a ratio.

We also assessed GMV at the voxel level. Structural MRI images were segmented to extract GM, WM, and CSF tissues using the unified segmentation algorithm in CAT12 (version 12.6, http://dbm.neuro.uni-jena.de/vbm) with default parameters. Warping to the Montreal Neurological Institute (MNI) template space was performed using Diffeomorphic Anatomical Registration Through Exponentiated Lie Algebra (DARTEL) with default parameters and registration to existing templates ^58^. Total intracranial volume (TIV) was computed as the sum of volumes of GM, WM, and CSF using the SPM “Estimate TIV and global tissue volumes” routine. Voxel-based statistical analyses were performed on the warped and modulated GMV maps, which were smoothed by a three-dimensional Gaussian kernel with full width at half maximum of 8 mm^3^.

#### 3.3.4 Additional measures

Age, sex, education, intelligence, SES, self-reported participation in physical activity and diagnostic group were considered as potential confounders. Educational attainment was measured in years of education. Crystallized intelligence was estimated using the Multiple-Choice Vocabulary Intelligence Test (MWT, min. score: 0, max. score: 37), with scores proportional to crystallized intelligence ^59^.

Participation in long-term physical activity was estimated using respective information from the LEQ. A mean score was calculated over responses on the frequency of physical activity over two or three life stages (i.e., < 65 years and ≥ 65 years, respectively). In addition, current physical activity was assessed through the Physical Activity Scale for the Elderly PASE, ^60^. The PASE includes leisure, household and occupational activities assessed over the previous week. Based on frequency, duration, and intensity of these activities, a total score is calculated with higher scores indicating greater levels of physical activity. Long-term physical activity was significantly correlated with current physical activity in the matched sample (*n* = 140, *r* = 0.35, *p* < 0.001), supporting the validity of the measure. Long-term physical activity was used as a covariate in statistical analyses, since the measure was available from all participants.

The SES was calculated for each participant using information on occupational activity assessed by the LEQ. Details are provided in the supplementary material. In brief, details on occupational activities of each respondent were obtained using 10 five-year intervals across middle-to late-life adulthood (i.e., 30 to 79 years of age). The information was used to calculate the international socio-economic index of occupational information (ISEI, min. score: 16, max. score: 90) ^61^ using a fully-automated procedure. The ISEI scores were averaged across time intervals to obtain one mean SES measure per participant. The SES measure was positively and significantly associated with the LEQ sum scores measuring educational as well as occupational activity for young (*n* = 140, *r* = 0.54, *p* < 0.001) and middle (*n* = 140, *r* = 0.69, *p* < 0.001) adulthood, indicating the validity of the estimated ISEI scores.

### 3.4 Statistical analyses

Statistical analyses were conducted using *R* (version 3.5.1.) and Statistical Parametric Mapping (SPM, version 12, Wellcome Trust Centre for Neuroimaging, London, UK). Figures were generated using the package *ggplot2* ^62^. Before conducting statistical models, statistical assumptions were assessed visually using diagnostic plots.

#### 3.4.1 Sample characteristics and matching procedure

Participants with lifelong musical activities and controls with no musical activity were matched using a one-to-one matching procedure taking into account age, sex, diagnostic group, education, SES, intelligence, and physical activity. Details are provided in the supplementary material. The procedure was carried out using propensity score matching with the *R* package *MatchIt* (version 4.1.0.) ^63^. Observations were matched based on the nearest-neighbor method, as a simple and effective procedure for selecting well-matched groups ^64^. Musical activity groups were compared in baseline demographic, behavioral, neuropsychological, and neuroimaging variables. Independent Student’s *t*-tests were used for all continuous variable and chi-squared (*χ*^*2*^) tests were applied for all categorical variables.

#### 3.4.2 ROI-based analyses

To assess our main hypotheses, multiple linear regression models were used. In these statistical analyses, an alpha value of 0.05 was considered statistically significant. In addition, correction for multiple comparisons was performed using a false discovery rate (FDR)-adjusted p-value threshold (alpha) of 0.05 ^65^. Uncorrected p-values were reported, when results survived FDR correction, this is specifically indicated.

Firstly, the association of musical activity (modelled as a main effect) with global cognition followed by the domain-specific abilities were assessed. Multiple linear regression models were performed including with musical activity (binary group variable) as an independent variable and each cognitive measure (z-transformed composite score) as a dependent variable, respectively. Next, the association between musical activity and brain structure was examined using similar multiple linear regressions. Models included musical activity (binary group) as independent variable and ROI-based GMV (frontal region and hippocampal region, both TIV adjusted) as dependent variable along with scanner site as covariate (dummy coded). Selected relationships were visualized to facilitate the interpretation of findings using box plots of unadjusted data.

Secondly, we assessed the moderating effect of musical activity on the relationship between ROI-based GMV and cognitive abilities. To do this, musical activity, ROI-based GMV (frontal region and hippocampal region, both mean-centered), the interaction term (musical activity × GMV), and scanner site as covariate (dummy coded) were entered into regression models with each cognitive factor score as dependent variable. To specify the directionality of the interactions, simple slope analyses were conducted ^66,67^. Interaction effects were visualized using unadjusted data as follows: General and domain-specific cognitive scores (z-transformed) were graphed as a function of musical activity and ROI-based GMV, respectively. In addition, we examined whether or not the respective relationships differed significantly from zero within each group.

#### 3.4.3 Voxel-based analysis

To further evaluate the spatial distribution of musical activity-associated effects on brain structure at the voxel level, exploratory voxel-wise general linear models (GLM) were conducted in SPM12. For the present purpose, voxel-wise results were presented at *p* < 0.001 uncorrected at peak level in combination with the estimated expected voxels per cluster (*k*) as automatically calculated by SPM.

Firstly, a GLM was computed with musical activity as independent variable and the modulated, warped, and smoothed GMV maps as dependent variable. Secondly, a moderating effect of musical activity was evaluated at the voxel level. This GLM included musical activity, the respective cognitive measure (z-transformed composite score), and the interaction term (musical activity × cognitive measure) as independent variables with GMV maps as dependent variable. The later analysis was carried out for global cognition and all cognitive domains. For reasons of simplicity, results of this analysis were displayed for one cognitive domain, selected by the strongest interaction effect in the ROI-based analysis.

All voxel-based analyses were adjusted for TIV as well as scanner site (dummy coded) and restricted to cerebral GM using an explicit binary GM mask derived from the present sample (i.e., average GM maps thresholded at a level of *>* 0.3, excluding cerebellum and brain stem). Cluster peaks are specified by their anatomical site, labelled using the Hammersmith atlas ^68^ provided by the CAT12 toolbox. Finally, mean values were extracted in significant clusters for each participant from the warped, modulated, and non-smoothed GMV images using the Marsbar toolbox (release: 0.44; http://marsbar.sourceforge.net/) ^69^, to provide complementary visualizations of the associations.

## 4 Results

### 4.1 Sample characteristics

This study included a total sample of 140 older participants (aged ≥ 60 years) selected from the ongoing, multi-center, observational DELCODE cohorts ^32^. The present sample comprised 70 individuals with lifelong regular musical activity and 70 controls with no musical activity over the life course (see Figure 1). The two groups (musical activity, no musical activity) were comparable in age, sex, distribution of diagnostic groups as well as reserve proxies of higher education, crystallized intelligence, SES, and participation in both long-term and current physical activity (all *p’s* > 0.05, Table 1). Slight group differences in frontal and total GMV (unadjusted raw values) were found, with larger volumes in the older participants with lifelong musical activity compared to controls.

**Table 1:** Descriptive characteristics of the matched sample (*n* = 140)

| Table 1: Descriptive characteristics of the matched sample (n = 140) |  |  |  |
| --- | --- | --- | --- |
|  | Musical activity | No musical activity | P value |
| Number (n) | 70 | 70 | - |
| Age (years) | 68.23 (6.62) | 69.01 (5.44) | 0.445 |
| Gender female/male (n) | 31/39 | 35/35 | 0.498 |
| Education (years) | 16.20 (2.71) | 15.96 (2.74) | 0.598 |
| Diagnostic group HC/ FH/SCD (n) | 19/7/44 | 24/6/40 | 0.654 |
| SES <sup>a</sup> | 66.27 (16.32) | 65.21 (16.04) | 0.699 |
| Crystallized intelligence <sup>b</sup> | 33.31 (2.14) | 33.04 (2.22) | 0.463 |
| Physical activity, long-term <sup>c</sup> | 4.25 (0.78) | 4.32 (0.71) | 0.611 |
| Physical activity, current <sup>d</sup> | 33.86 (11.80), n = 66 | 32.45 (12.85), n = 69 | 0.507 |
| Total hippocampal GMV (sum, ml) | 6.26 (0.71) | 6.21 (0.66) | 0.692 |
| Total frontal GMV (ml) | 138.86 (12.44) | 134.69 (11.88) | 0.044* |
| Total cortical GMV (ml) | 453.83 (37.56) | 441.69 (37.13) | 0.049* |
| Descriptive data are given if applicable as mean and standard deviation (in parenthesis). The actual sample size is provided, if different from sample size specified in first row. P values correspond to independent t-tests for unequal variance with participant group as independent variable. Chi-square statistic was used to compare the distribution of categorical variables.<br>***p < 0.001, **p < 0.01, *p < 0.05.<br>**Key:** HC, healthy control participants; FH, participants with family history of AD; GMV, gray matter volume; SCD, participants with subjective cognitive decline; SES, socioeconomic status.<br><sup>a</sup> International socio-economic index (ISEI); <sup>b</sup> Multiple-Choice Vocabulary Intelligence Test (MWT); <sup>c</sup> Lifetime of Experiences Questionnaire (LEQ); <sup>d</sup> Physical Activity Scale for the Elderly (PASE). |  |  |  |

### 4.2 Musical activity and cognition

Applying multiple linear regression models to assess the associations between musical activity and cognitive performance, we found significant group differences for global cognition, working memory, executive function, language and visuospatial abilities (Table 2). Performance in these cognitive domains was significantly better in the older participants with lifelong regular musical activity compared to controls. In contrast, no association of musical activity was found with the domain of learning and memory (*p* = 0.209).

**Table 2:**
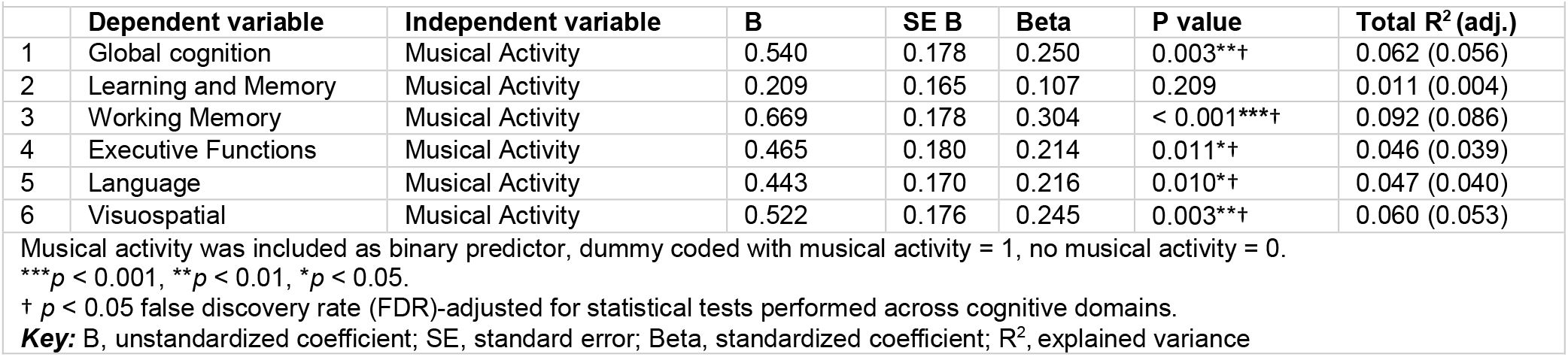
Results of linear regression analyses between musical activity and cognition.

### 4.3 Musical activity and brain structure in regions-of-interest

Results for the associations between musical activity and GMV (TIV-adjusted values) in the selected ROIs are shown in Table 3. There were no significant differences between participants with lifelong musical practice compared to controls in frontal and hippocampal volume (all *p’s* > 0.5). Also, the groups did not differ significantly in total cortical GMV (*p* = 0.722).

**Table 3:** Results of linear regression analyses between musical activity and GMV in regions-of-interest.

|  |  |  |  |  |  |  |  |
| --- | --- | --- | --- | --- | --- | --- | --- |
|  | Dependent variable | Independent variable | B | SE B | Beta | P value | Total R<sup>2</sup> (adj.) |
| 1 | Frontal GMV | Musical Activity | -0.046 | 0.204 | -0.020 | 0.822 | 0.120 (0.052) |
| 1 | Hpc GMV | Musical Activity | -0.007 | 0.012 | -0.052 | 0.551 | 0.102 (0.032) |
| 3 | Cortical GMV | Musical Activity | -0.303 | 0.850 | -0.031 | 0.722 | 0.132 (0.065) |

Models adjusted for scanner site. Musical activity was included as binary predictor, dummy coded with musical activity = 1, no musical activity = 0. Regional GMV was adjusted by total intracranial volume (TIV). \*\*\* $p < 0.001$ , \*\* $p < 0.01$ , \* $p < 0.05$ . † $p < 0.05$ false discovery rate (FDR)-adjusted for statistical tests performed across ROIs. **Key:** B, unstandardized coefficient; Hpc, Hippocampus; SE, standard error; Beta, standardized coefficient; R<sup>2</sup>, explained variance; GMV, gray matter volume

### 4.4 Moderations of musical activity in regions-of-interest

Moderation analyses were applied to assess the influence of musical activity on the associations between late-life regional GMV and cognitive performance (Table 4 and 5). Frontal GMV was positively associated with global cognition and domain-specific cognitive abilities (all *p’s* ≤ 0.001, data not shown). Importantly though, significant interactions were observed between musical activity and frontal GMV for global cognition, working memory, and language abilities (all *p’s* < 0.05; Table 4). Visualization of these relationships (Figure 3) indicated that these cognitive abilities were selectively enhanced in participates with musical activity and preserved GMV in the frontal regions (i.e., above the 90% percentile of the GMV distribution in AD patients). No similar effect was detected for the learning and memory domain. Hippocampal GMV was also positively associated with global cognition and domain-specific cognitive abilities (all *p’s* < 0.01, data not shown). There were, however, no significant interactions between musical activity and hippocampal volume in the multi-domain cognitive abilities (all *p’s* > 0.1, Table 5 and supplementary Figure 4).

**Table 4:** Results of the interaction analyses between musical activity and frontal GMV.

|  |  |  |  |  |  |  |  |
| --- | --- | --- | --- | --- | --- | --- | --- |
|  | **Dependent variable** | **Independent variable** | **B** | **SE B** | **Beta** | **P value** | **Total R<sup>2</sup> (adj.)** |
| 1 | Global cognition | Music Activity × Frontal GMV | 0.318 | 0.139 | 0.261 | 0.024*† | 0.332 (0.269) |
| 2 | Learning and Memory | Music Activity × Frontal GMV | 0.102 | 0.132 | 0.092 | 0.441 | 0.263 (0.193) |
| 3 | Working Memory | Music Activity × Frontal GMV | 0.432 | 0.141 | 0.348 | 0.003**† | 0.335 (0.273) |
| 4 | Executive Functions | Music Activity × Frontal GMV | 0.278 | 0.145 | 0.228 | 0.058 | 0.273 (0.204) |
| 5 | Language | Music Activity × Frontal GMV | 0.316 | 0.133 | 0.274 | 0.019*† | 0.320 (0.256) |
| 6 | Visuospatial | Music Activity × Frontal GMV | 0.227 | 0.145 | 0.189 | 0.119 | 0.251 (0.180) |

Models adjusted for scanner site. Musical activity was included as binary predictor, dummy coded with musical activity = 1, no musical activity = 0. Frontal GMV was adjusted for total intracranial volume and mean centered. \*\*\* $p < 0.001$ , \*\* $p < 0.01$ , \* $p < 0.05$ . † $p < 0.05$ false discovery rate (FDR)-adjusted for statistical tests performed across cognitive domains. **Key:** B, unstandardized coefficient; SE, standard error; Beta, standardized coefficient; R<sup>2</sup>, explained variance

**Table 5:** Results of the interaction analyses between musical activity and hippocampal GMV.

|  |  |  |  |  |  |  |  |
| --- | --- | --- | --- | --- | --- | --- | --- |
|  | **Dependent variable** | **Independent variable** | **B** | **SE B** | **Beta** | **P value** | **Total R<sup>2</sup> (adj.)** |
| 1 | Global cognition | Music Activity × Hpc GMV | 2.248 | 2.515 | 0.118 | 0.373 | 0.330 (0.266) |
| 2 | Learning and Memory | Music Activity × Hpc GMV | -0.727 | 2.350 | -0.042 | 0.757 | 0.284 (0.216) |
| 3 | Working Memory | Music Activity × Hpc GMV | 4.109 | 2.620 | 0.211 | 0.119 | 0.299 (0.233) |
| 4 | Executive Functions | Music Activity × Hpc GMV | 2.268 | 2.640 | 0.118 | 0.392 | 0.268 (0.199) |
| 5 | Language | Music Activity × Hpc GMV | 2.782 | 2.435 | 0.154 | 0.255 | 0.300 (0.234) |
| 6 | Visuospatial | Music Activity × Hpc GMV | 0.897 | 2.547 | 0.047 | 0.725 | 0.290 (0.223) |

Models adjusted for scanner site. Musical activity was included as binary predictor, dummy coded with musical activity = 1, no musical activity = 0. Hippocampal GMV was adjusted for total intracranial volume and mean centered. \*\*\* $p < 0.001$ , \*\* $p < 0.01$ , \* $p < 0.05$ . † $p < 0.05$ false discovery rate (FDR)-adjusted for statistical tests performed across cognitive domains. **Key:** B, unstandardized coefficient; Hpc, hippocampus; SE, standard error; Beta, standardized coefficient; R<sup>2</sup>, explained variance

**Figure 3:**
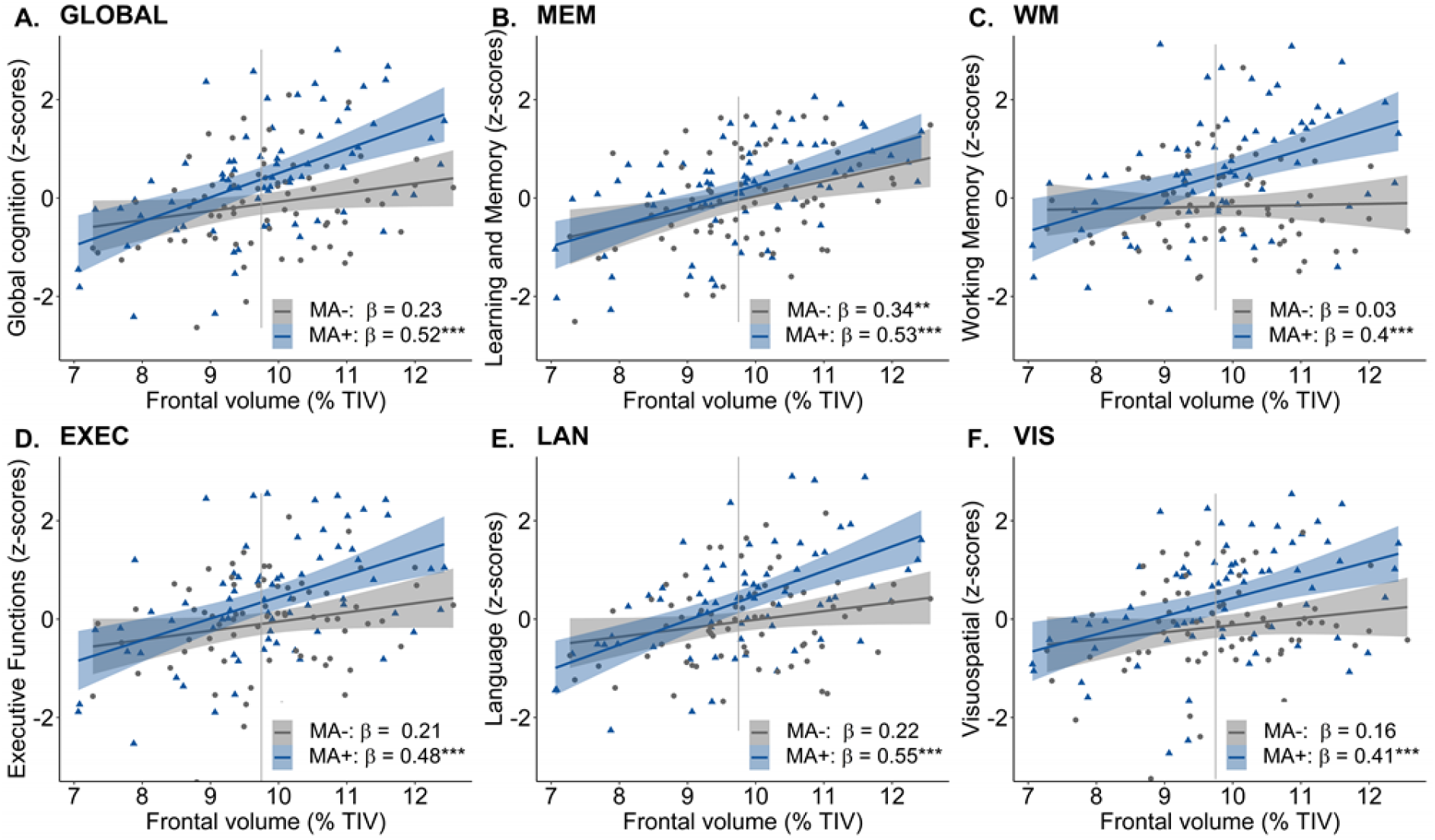
Moderation effect of lifelong musical activity in the frontal region. A significant moderation effect of musical activity was observed for global cognition (A, GLOBAL), working memory (C, WM), and language abilities (E, LAN), such that larger frontal volume (above the 90th percentile of the frontal volume distribution in AD patients) was associated with better global in participants with lifelong musical activity (MA+, blue) compared to controls (MA-, gray). This interaction was not significant for learning and memory (B, MEM), executive functions (D, EXEC), and visuospatial abilities (F, VIS). Individual data points (dots and triangles), linear trends (solid lines), 95% confidence intervals (shaded areas), and standardized regression coefficients (β) within each group are provided. Gray vertical lines display the 90th percentile of the frontal GMV distribution in AD patients of the DELCODE study. Significance levels (uncorrected): \*\*\**p* < 0.001, \*\**p* < 0.01, \**p* < 0.05. ***Key:*** GMV, gray matter volume; MA+, musical activity; MA-, no musical activity; TIV, total intracranial volume.

### 4.5 Voxel-based analysis

Results of the exploratory analyses at the voxel level are presented in Table 6 and Figure 4. There was a subtle positive association between lifelong regular musical activity and GMV within a smaller cluster in the left postcentral gyrus (*p* < 0.001 uncorrected). No other significant clusters were found. The interaction analysis corroborated a significant moderation of musical activity on the association between working memory and regional GMV (*p* < 0.001 uncorrected) in frontal (lateral and medial), inferior temporal, and precentral regions. Results of the moderation analyses across all cognitive measures were essentially similar, with some variations in the number of significant clusters (supplementary Figure 6). No significant interaction effect was found for the domain of learning and memory at the voxel level.

**Figure 4:**
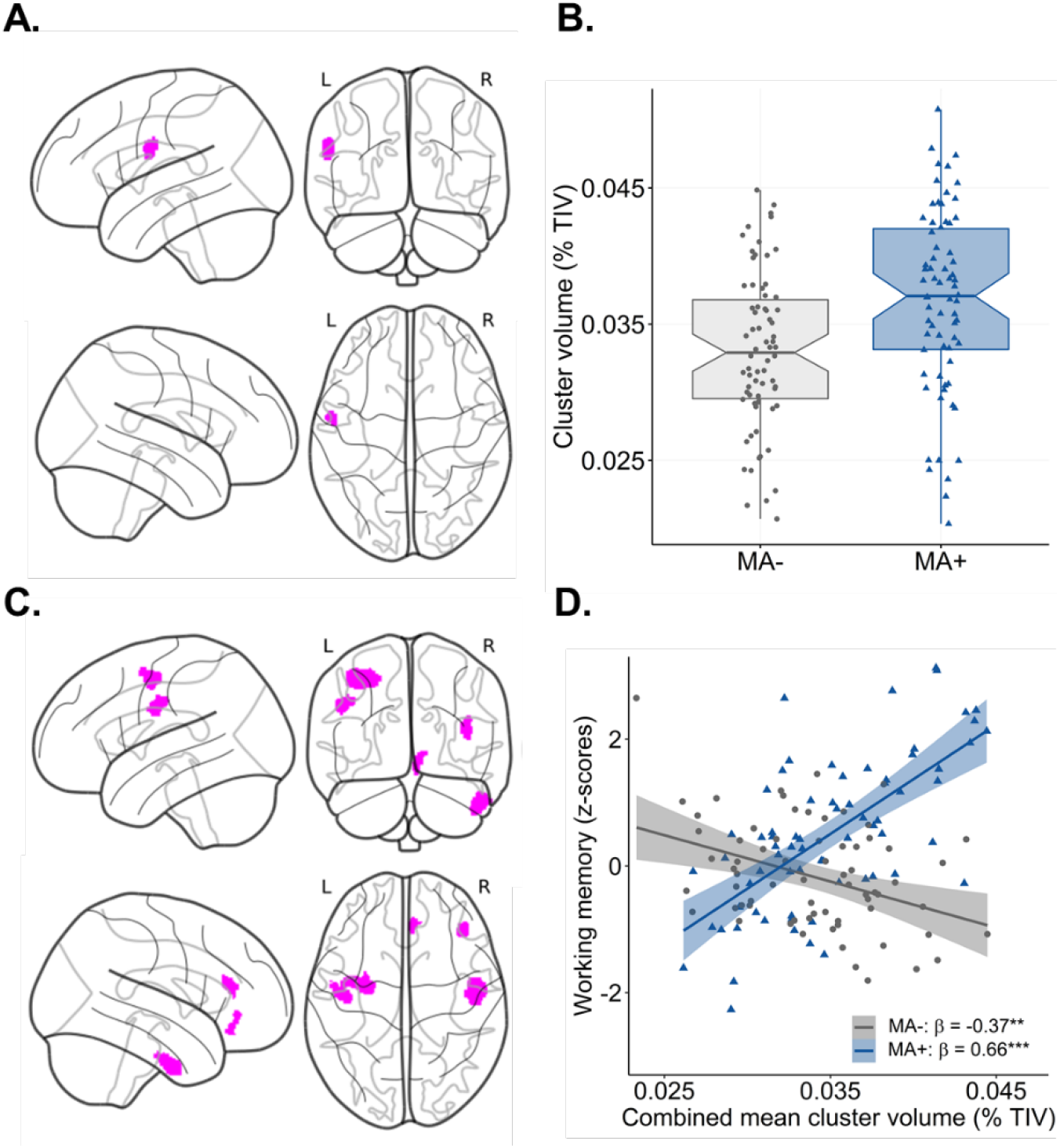
Associations between lifelong musical activity and regional volume distribution. A-B. Results of the main effect analysis. Statistical map (A) shows significant clusters (*p* < 0.001 uncorrected, color-coded in magenta) with larger GMV in participants with musical activity compared to controls. The corresponding graph (B) displays the association using mean GMV values extracted from the corresponding cluster in the postcentral gyrus. The box plot displays the median with 95% confidence intervals, interquartile range with lower (25th) and upper percentiles (75th), and individual data points. **C-D. Results of the interaction analysis**. The statistical map (C) displays clusters (*p* < 0.001 uncorrected, color-coded in magenta) with a significant moderation effect of musical activity. The corresponding scatter plot (D) shows the association using mean values extracted from the GMV maps in the combined cluster. Larger GMV in the combined cluster was associated with better working memory ability selectively in the musically active participants (MA+, blue) compared to controls (MA-, gray). Individual data points, linear trends (solid lines), 95% confidence intervals (shaded areas), and standardized regression coefficients (β) within each musical activity group are provided. The statistical maps are depicted on a glass brain. Significance levels (uncorrected): \*\*\**p* < 0.001, \*\**p* < 0.01, \**p* < 0.05. ***Key:*** MA+, musical activity; MA-, no musical activity; GMV, gray matter volume; TIV, total intracranial volume.

**Table 6:** Results of analyses between musical activity and GMV at the voxel level.

|  |  |  |  |  |  |  |  |  |  |
| --- | --- | --- | --- | --- | --- | --- | --- | --- | --- |
| Model / contrast | No. cluster | Label | Hemisphere | Cluster |  | Peak of cluster |  |  |  |
|  |  |  |  | p | size | Z value | MNI coordinates (x y z) |  |  |
| Main effect <sup>a</sup> |  |  |  |  |  |  |  |  |  |
| Positive | 1 | Postcentral gyrus | left | 0.176 | 239 | 3.66 | -57 | -6 | 26 |
| Interaction effect <sup>b</sup> |  |  |  |  |  |  |  |  |  |
|  | 1 | Precentral gyrus | left | 0.019 | 646 | 4.83 | -28 | -10 | 48 |
|  | 2 | Precentral gyrus | left | 0.123 | 251 | 4.46 | -38 | -8 | 52 |
|  | 3 | Inferior middle temporal gyrus | right | 0.048 | 437 | 4.42 | 51 | -12 | -42 |
|  | 4 | Inferior frontal gyrus | right/<br>lateral | 0.171 | 194 | 3.70 | 40 | 33 | 12 |
|  | 5 | Superior frontal gyrus | right/<br>medial | 0.273 | 122 | 3.47 | 6 | 36 | -12 |
| Models adjusted for scanner site and TIV. |  |  |  |  |  |  |  |  |  |
| Musical activity was included as binary predictor, dummy coded with musical activity = 1, no musical activity = 0. |  |  |  |  |  |  |  |  |  |
| <sup>a</sup> Results from the main effect model with musical activity and GMV ($p < 0.001$ uncorrected, expected voxels per cluster $k = 139$). | | | | | | | | | |
| <sup>b</sup> Results from the interaction effect model with musical activity, working memory, and GMV ($p < 0.001$ uncorrected, expected voxels per cluster $k = 110$). | | | | | | | | | |
| Cluster peaks are specified by their anatomical site, labelled using the Hammersmith atlas provided by the CAT12 toolbox. |  |  |  |  |  |  |  |  |  |
| **Key:** GMV, gray matter volume; MNI x y z [mm], coordinates MNI space in millimeters; TIV, total intracranial volume |  |  |  |  |  |  |  |  |  |

## 5 Discussion

### 5.1 Summary

The current study examined late-life cognitive abilities, brain morphology, and their interplay in cognitively healthy older adults as a function of regularly playing a musical instrument over the life course. Participants with a self-reported history of lifelong musical activity were compared to matched controls without musical activity across the lifespan. Results of this study highlight that regularly playing a musical instrument is associated with global and multi-domain cognitive benefits in older adults, with no significant benefit in gray matter structure in regions affected by aging and AD. In the musically active participants, cognitive abilities were enhanced with preserved regional GMV for some cognitive domains, pointing towards a facilitated recruitment of existing brain resources in this group. Overall, our findings may imply that a history of regular musical activity could promote cognitive and brain benefits in older adults and thereby strengthen resilience against cognitive decline.

### 5.2 Musical activity and cognition

We demonstrate that participants, who reported a lifelong regular engagement in musical activity, outperformed matched controls in cognitive performance. More precisely, superior cognitive abilities were found in global cognition and multiple cognitive domains including working memory, executive functions, language and visuospatial abilities in the musically active older people, with the largest effect size seen for working memory. These findings directly support and expand previous studies, showing that playing a musical instrument may preserve higher-order cognitive skills that typically decline in older adults ^14,15,33,34^. By contrast, we did not identify benefits of musical activity on learning and memory, although these must be perceived as essential cognitive skill involved in playing music. In older adults, some studies report that regular musical activity is associated with better episodic memory ^13,14,33^, while others do not ^8,15,16^. It might, however, be argued that specific hippocampus-related processes are enhanced by musical activity, such as long-term musical memory or navigation in acoustic space ^27,35^, but cannot be captured by the memory composite used in the current study. While sensitive experimental and neuroimaging markers are needed to gain insights into presumed memory benefits, current findings appear to confirm that regular musical activity may particularly favor late-life cognitive abilities involving the frontal lobe.

### 5.3 Musical activity and brain structure

Our data demonstrate that playing music during life was not significantly associated with larger GMV in age-sensitive brain regions. Specifically, we did not detect volumetric differences between musically active people and controls in frontal or hippocampal regions. The voxel-based analysis confirmed this observation. A slight volume increase was found in somatosensory areas of the musically active group, presumably reflecting brain plasticity in response to tactile stimulations induced by playing a instrument ^21,22^. However, we failed to identify respective GMV modulations in higher-order brain regions. Earlier studies have shown positive associations between musical activity and brain volume in fontal, temporal, and parietal regions mainly in younger cohorts ^21,22,27^, with limited indication in older adults ^30^. Similar to previous studies ^15,16,33^, we accounted for several reserve proxies that may help maintain late-life brain structure ^3,36^. Given this effort, it seems plausible to assume that there is little benefit of regular musical activity on structural brain resources in older age. Alternatively, subtle effects could be unnoticed due to increased variability in GMV through differential brain aging and/or brain pathology ^37^.

### 5.4 Moderations of musical activity

Importantly though, our current results revealed that lifelong regular musical activity could act as a protective factor in the associations between late-life brain resources and cognitive performance. We found an interaction between playing music and GMV, such that performance in some cognitive domains was enhanced with preserved frontal volume selectively in the musically active participants. This specific moderation effect was significant for global cognition, language, as well as working memory and extended to inferior temporal as well as motor-sensory regions at the voxel level. In other words, although gray matter structure was not substantially associated with musical activity, it facilitated late-life cognitive performance in synergy with playing music. This may reflect a more efficient use of an overall younger brain age, as previously reported in amateur musicians compared to controls ^38^. Our observation further parallels existing findings ^29^. In their study, a larger hippocampal volume was associated with better general cognitive abilities in younger musicians, but not in non-musicians, implying that musically active people may be able to use existing brain resources more efficiently ^29^. The current findings essentially indicate that this functional advantage of playing music is detectable in older adults, where it is linked to distributed brain regions and multiple cognitive domains. Frontal and temporal brain regions, in particular, are part of wide-spread brain networks shown to support cognitive reserve processes ^36,39-42^. It therefore appears that playing music over the life course could facilitate the recruitment of structural brain resources, as a key benefit to support late-life cognitive functioning.

### 5.5 Synopsis

Taken together, the present study adds considerable insight to the picture that musical activity over the life course, even at a moderate frequency, could act as protective factor in late life stages. Given that playing a musical instrument requires the simultaneous integration of intense multimodal motor, sensory, cognitive, emotional, and social sensations, this lifestyle activity may induce lasting functional plasticity in higher-order neural networks supporting multi-domain cognitive functions ^43,44^. Even passive listening to music was previously shown to modulate functional connectivity in distributed brain networks ^45^, a mechanism suggested to convey therapeutic benefits of music-based interventions ^7^. Enhanced functional connectivity in higher-order brain networks is an essential mechanism shown to be protective against neuropathological burden in older adults ^39,41^. In light of our findings, it may be proposed that musical activity could act as a resilience factor through functional brain resources, which need to be examined in future studies.

Nevertheless, the observed health benefits associated with lifelong regular musical activity could be encouraged by a general engagement in advantageous lifestyles. Similar to previous studies ^33^, participants with regular musical activity during the life course were characterized by a high reserve profile including higher education, SES, intelligence, and more frequent physical activity. Notably though, we observed better cognitive abilities in the musically active group with these factors accounted for, suggesting an added benefit of musical activity on late-life brain and cognitive functions beyond known reserve proxies. Overall, our results propose that playing a musical instrument could serve as a low-threshold multimodal enrichment strategy that may help preserve cognitive and brain health. Targeted intervention studies are required to evaluate the impact of playing music on cognitive performance and underlying brain mechanisms in older people ^46^.

### 5.6 Strengths and limitations

Our study has several strengths and limitations. We assembled data from the longitudinal observational DELCODE cohort to assess a well-characterized sample of cognitively unimpaired older adults with measures of demographics, cognition, lifestyle behaviour, and brain structure. This detailed phenotyping provided new evidence on potential health benefits of regular musical activity in older adults. This study identified older people with a history of musical activity over three life stages and statistical analyses were based on multi-domain cognitive abilities and morphological brain measures all measured in the same participants.

Limitations of our cohort-based approach include the assessment of musical activity. While we obtained the frequency of musical activity using the LEQ ^47^, more detailed information on musical instrument type, age of acquisition, and intensity would be desirable given that these features may differentially impact brain plasticity and cognitive skills ^14,48^. Furthermore, the present cross-sectional study design does not permit causal interpretations of the investigated associations. It might be possible that certain factors that were not accounted for, facilitate playing an instrument over the life course. Such factors may include genetic predispositions or advantageous early-life exposures, which could play a role in the observed relationships, warranting further investigations. Finally, potential benefits of musical activity call for a longitudinal study design, to evaluate if musically active older people are indeed more protected against cognitive decline, which will have important implications on public health strategies.

### 5.7 Conclusion

Results of the present study are promising and suggest that lifelong regular musical activity, as an accessible and multimodal leisure activity, could help mitigating age-related cognitive decline through benefits in functional brain resources. Further research is needed to assess detailed information about the nature of playing music and functional brain mechanisms associated with a history of regular musical activity over the life course. Given that world populations are aging and that age-related diseases pose healthcare challenges of utmost importance, interventional studies examining the protective effects of musical activity on the brain and cognitive functioning in older adults are greatly needed.

## 6 Availability statements

### 6.1 Data availability

The data that support findings of the present study are available on reasonable request.

### 6.2 Code availability

For this study, existing data analysis packages for statistical analyses were used. Scripts for the use of these packages are available online from the authors on reasonable request.

## 7 Acknowledgements

We are grateful for the tremendous efforts of all DELCODE study teams across participating DZNE sites. Further, we express our sincere gratitude to all volunteers and their family members that participate in the DELCODE study. We gratefully acknowledge all administrative and scientists staff members involved in data acquisition, data management, as well as quality control. We thank Andrea Dell’Orco for his support in data visualisation.

DELCODE study group: H. Amthauer, A. Cetindag, N. Cosma, D. Diesing, M. Ehrlich, F. Fenski, S. Freiesleben, M. Fuentes, D. Hauser, N. Hujer, E. Incesoy, C. Kainz, C. Lange, K. Lindner, H. Megges, O. Peters, L. Preis, S. Altenstein, A. Lohse, C. Franke, J. Priller, E. Spruth, I. Villar Munoz, M. Barkhoff, H. Boecker, F. Brosseron, M. Daamen, T. Engels, J. Faber, K. Fließbach, I. Frommann, M. Grobe-Einsler, G. Hennes, G. Herrmann, L. Jost, P. Kalbhen, O. Kimmich, X. Kobeleva, B. Kofler, C. McCormick, L. Miebach, C. Miklitz, A. Müller, D. Oender, A. Polcher, V. Purrer, S. Röske, C. Schneider, A. Schneider, A. Spottke, I. Vogt, M. Wagner, S. Wolfsgruber, S. Yilmaz, C. Bartels, P. Dechent, N. Hansen, L. Hassoun, S. Hirschel, S. Nuhn, I. Pfahlert, L. Rausch, B. Schott, C. Timäus, C. Werner, J. Wiltfang, L. Zabel, H. Zech, A. Bader, J.C. Baldermann, B. Dölle, A. Drzezga, C. Escher, N. Ghiasi, K. Hardenacke, F. Jessen, H. Lützerath, F. Maier, B. Marquardt, A. Martikke, D. Meiberth, S. Petzler, A. Rostamzadeh, L. Sannemann, A. Schild, S. Sorgalla, S. Stockter, M. Thelen, M. Tscheuschler, F. Uhle, P. Zeyen, D. Bittner, A. Cardenas-Blanco, L. Dobisch, E. Düzel, D. Grieger-Klose, D. Hartmann, C. Metzger, P. Nestor, C. Ruß, F. Schulze, O. Speck, G. Wenzel, R. Yakupov, G. Ziegler, C. Brauneis, K. Bürger, C. Catak, L. Coloma Andrews, M. Dichgans, A. Dörr, B. Ertl-Wagner, D. Frimmer, B. Huber, D. Janowitz, M. Kreuzer, E. Markov, C. Müller, A. Rominger, J. Schmid, A. Seegerer, J. Stephan, A. Zollver, L. Burow, S. de Jonge, P. Falkai, N. Garcia Angarita, T. Görlitz, S. Gürsel, I. Horvath, C. Kurz, E. Meisenzahl-Lechner, R. Perneczky, J. Utecht, M. Dyrba, H. Janecek-Meyer, I. Kilimann, C. Lappe, E. Lau, H. Pfaff, H. Raum, P. Sabik, M. Schmidt, H. Schulz, S. Schwarzenboeck, S. Teipel, M. Weber, M. Buchmann, T. Heger, P. Hinderer, E. Kuder-Buletta, C. Laske, M. Munk, C. Mychajliw, S. Soekadar, P. Sulzer, T. Trunk

## 8 Declarations

### 8.1 Ethics approval and consent to participate

The DELCODE study protocol was approved by the ethical committees of the medical faculties of participating sites. All participants gave written informed consent prior to study inclusion.

### 8.2 Availability of data and materials

The data that support findings of this study are available on reasonable request.

### 8.5 Authors’ contributions

Conceptualization and design of the current study: A.B., T.K., A.H., K.F., S.R., M.Wa., G.K., M.Wi.; Overall design and implementation of the DELCODE study: O.P., S.D.F., J.P., S.A., A.S., K.F., I.F., A.S., N.R., S.W., L.K., J.W., C.B., F.M., C.N., L.D., R.Y., K.B., D.J., R.P., B.R., S.T., I.K., C.L., M.M., J.D.H., P.D., M.E., K.S., E.D., F.J., S.R., M.Wa.; methodology/statistical analysis: A.B., A.Z., M.G., M.W.; interpretation of data: A.B., A.Z., T.K., A.H., K.F., S.W., S.R., M.Wa., G.K., M.Wi.; drafting and/or revision of manuscript and approval of the final version: : A.B., A.Z., T.K., M.G., A.H., K.F., O.P., S.D.F., J.P., S.A., A.S., K.F., I.F., A.S., N.R., S.W., L.K., J.W., C.B., F.M., C.N., L.D., R.Y., K.B., D.J., R.P., B.R., S.T., I.K., C.L., M.M., J.D.H., P.D., M.E., K.S., E.D., F.J., S.R., M.Wa., G.K., M.Wi.

## References

1. Livingston, G., et al. Dementia prevention, intervention, and care: 2020 report of the Lancet Commission. Lancet (London, England) 396, 413–446 (2020).

2. Arenaza-Urquijo, E.M., Wirth, M. & Chetelat, G. Cognitive reserve and lifestyle: moving towards preclinical Alzheimer’s disease. Frontiers in aging neuroscience 7, 134 (2015).

3. Wirth, M., Haase, C.M., Villeneuve, S., Vogel, J. & Jagust, W.J. Neuroprotective pathways: lifestyle activity, brain pathology, and cognition in cognitively normal older adults. Neurobiol Aging 35, 1873–1882 (2014).

4. Valenzuela, M.J., et al. Multiple biological pathways link cognitive lifestyle to protection from dementia. Biological psychiatry 71, 783–791 (2012).

5. Verghese, J., et al. Leisure activities and the risk of dementia in the elderly. N Engl J Med 348, 2508–2516 (2003).

6. Balbag, M.A., Pedersen, N.L. & Gatz, M. Playing a Musical Instrument as a Protective Factor against Dementia and Cognitive Impairment: A Population-Based Twin Study. Int J Alzheimers Dis 2014, 836748 (2014).

7. Sihvonen, A.J., et al. Music-based interventions in neurological rehabilitation. Lancet Neurol 16, 648–660 (2017).

8. Fauvel, B., et al. Musical practice and cognitive aging: two cross-sectional studies point to phonemic fluency as a potential candidate for a use-dependent adaptation. Frontiers in aging neuroscience 6, 227 (2014).

9. Kempermann, G. Environmental enrichment, new neurons and the neurobiology of individuality. Nature Reviews Neuroscience 20, 235–245 (2019).

10. Lappe, C., Herholz, S.C., Trainor, L.J. & Pantev, C. Cortical plasticity induced by short-term unimodal and multimodal musical training. J Neurosci 28, 9632–9639 (2008).

11. Román-Caballero, R., Arnedo, M., Triviño, M. & Lupiáñez, J. Musical practice as an enhancer of cognitive function in healthy aging - A systematic review and meta-analysis. PLoS One 13, e0207957–e0207957 (2018).

12. Sutcliffe, R., Du, K. & Ruffman, T. Music Making and Neuropsychological Aging: A Review. Neuroscience & Biobehavioral Reviews 113, 479–491 (2020).

13. Gooding, L.F., Abner, E.L., Jicha, G.A., Kryscio, R.J. & Schmitt, F.A. Musical Training and Late-Life Cognition. Am J Alzheimers Dis Other Demen 29, 333–343 (2014).

14. Hanna-Pladdy, B. & MacKay, A. The relation between instrumental musical activity and cognitive aging. Neuropsychology 25, 378–386 (2011).

15. Hanna-Pladdy, B. & Gajewski, B. Recent and past musical activity predicts cognitive aging variability: direct comparison with general lifestyle activities. Front Hum Neurosci 6, 198 (2012).

16. Strong, J.V. & Mast, B.T. The cognitive functioning of older adult instrumental musicians and non-musicians. Neuropsychol Dev Cogn B Aging Neuropsychol Cogn 26, 367–386 (2019).

17. Gray, R. & Gow, A.J. How is musical activity associated with cognitive ability in later life? Neuropsychol Dev Cogn B Aging Neuropsychol Cogn 27, 617–635 (2020).

18. Schneider, C.E., Hunter, E.G. & Bardach, S.H. Potential Cognitive Benefits From Playing Music Among Cognitively Intact Older Adults: A Scoping Review. J Appl Gerontol 38, 1763–1783 (2019).

19. Herholz, S.C. & Zatorre, R.J. Musical training as a framework for brain plasticity: behavior, function, and structure. Neuron 76, 486–502 (2012).

20. Wan, C.Y. & Schlaug, G. Music making as a tool for promoting brain plasticity across the life span. Neuroscientist 16, 566–577 (2010).

21. Gaser, C. & Schlaug, G. Brain structures differ between musicians and non-musicians. J Neurosci 23, 9240–9245 (2003).

22. Gärtner, H., et al. Brain morphometry shows effects of long-term musical practice in middle-aged keyboard players. Frontiers in psychology 4, 636 (2013).

23. Fauvel, B., et al. Morphological brain plasticity induced by musical expertise is accompanied by modulation of functional connectivity at rest. Neuroimage 90, 179–188 (2014).

24. Habes, M., et al. White matter hyperintensities and imaging patterns of brain ageing in the general population. Brain 139, 1164–1179 (2016).

25. Jack, C.R., Jr., et al. Medial temporal atrophy on MRI in normal aging and very mild Alzheimer’s disease. Neurology 49, 786–794 (1997).

26. Wirth, M., et al. Regional patterns of gray matter volume, hypometabolism, and beta-amyloid in groups at risk of Alzheimer’s disease. Neurobiol Aging 63, 140–151 (2018).

27. Groussard, M., et al. When music and long-term memory interact: effects of musical expertise on functional and structural plasticity in the hippocampus. PLoS One 5 (2010).

28. Herdener, M., et al. Musical Training Induces Functional Plasticity in Human Hippocampus. The Journal of Neuroscience 30, 1377–1384 (2010).

29. Oechslin, M.S., et al. Hippocampal volume predicts fluid intelligence in musically trained people. Hippocampus 23, 552–558 (2013).

30. Chaddock-Heyman, L., et al. Musical Training and Brain Volume in Older Adults. Brain Sci 11 (2021).

31. Stern, Y., et al. Whitepaper: Defining and investigating cognitive reserve, brain reserve, and brain maintenance. Alzheimers Dement 16, 1305–1311 (2020).

32. Jessen, F., et al. Design and first baseline data of the DZNE multicenter observational study on predementia Alzheimer’s disease (DELCODE). Alzheimers Res Ther 10, 15 (2018).

33. Mansens, D., Deeg, D.J.H. & Comijs, H.C. The association between singing and/or playing a musical instrument and cognitive functions in older adults. Aging Ment Health 22, 964–971 (2018).

34. Groussard, M., Coppalle, R., Hinault, T. & Platel, H. Do Musicians Have Better Mnemonic and Executive Performance Than Actors? Influence of Regular Musical or Theater Practice in Adults and in the Elderly. Frontiers in human neuroscience 14, 557642 (2020).

35. Teki, S., et al. Navigating the auditory scene: an expert role for the hippocampus. J Neurosci 32, 12251–12257 (2012).

36. Arenaza-Urquijo, E.M., et al. Relationships between years of education and gray matter volume, metabolism and functional connectivity in healthy elders. Neuroimage 83C, 450–457 (2013).

37. Hedden, T. & Gabrieli, J.D. Insights into the ageing mind: a view from cognitive neuroscience. Nat Rev Neurosci 5, 87–96 (2004).

38. Rogenmoser, L., Kernbach, J., Schlaug, G. & Gaser, C. Keeping brains young with making music. Brain Struct Funct 223, 297–305 (2018).

39. Benson, G., et al. Functional connectivity in cognitive control networks mitigates the impact of white matter lesions in the elderly. Alzheimers Res Ther 10, 109 (2018).

40. Colangeli, S., et al. Cognitive Reserve in Healthy Aging and Alzheimer’s Disease: A Meta-Analysis of fMRI Studies. Am J Alzheimers Dis Other Demen 31, 443–449 (2016).

41. Franzmeier, N., Duering, M., Weiner, M., Dichgans, M. & Ewers, M. Left frontal cortex connectivity underlies cognitive reserve in prodromal Alzheimer disease. Neurology 88, 1054–1061 (2017).

42. Marques, P., et al. The functional connectome of cognitive reserve. Hum Brain Mapp 37, 3310–3322 (2016).

43. Oechslin, M.S., Van De Ville, D., Lazeyras, F., Hauert, C.-A. & James, C.E. Degree of Musical Expertise Modulates Higher Order Brain Functioning. Cereb Cortex 23, 2213–2224 (2012).

44. Fauvel, B., Groussard, M., Eustache, F., Desgranges, B. & Platel, H. Neural implementation of musical expertise and cognitive transfers: could they be promising in the framework of normal cognitive aging? Frontiers in human neuroscience 7, 693 (2013).

45. Karmonik, C., et al. Music Listening modulates Functional Connectivity and Information Flow in the Human Brain. Brain connectivity 6, 632–641 (2016).

46. James, C.E., et al. Train the brain with music (TBM): brain plasticity and cognitive benefits induced by musical training in elderly people in Germany and Switzerland, a study protocol for an RCT comparing musical instrumental practice to sensitization to music. BMC Geriatr 20, 418 (2020).

47. Valenzuela, M.J. & Sachdev, P. Assessment of complex mental activity across the lifespan: development of the Lifetime of Experiences Questionnaire (LEQ). Psychol Med 37, 1015–1025 (2007).

48. Bangert, M. & Schlaug, G. Specialization of the specialized in features of external human brain morphology. Eur J Neurosci 24, 1832–1834 (2006).

49. Morris, J.C., et al. The Consortium to Establish a Registry for Alzheimer’s Disease (CERAD). Part I. Clinical and neuropsychological assessment of Alzheimer’s disease. Neurology 39, 1159–1165 (1989).

50. Jessen, F., et al. A conceptual framework for research on subjective cognitive decline in preclinical Alzheimer’s disease. Alzheimers Dement 10, 844–852 (2014).

51. Roeske, S., et al. P3-591: A German version of the lifetime of experiences questionnaire (LEQ) to measure cognitive reserve: Validation results from the DELCODE study. Alzheimer’s & Dementia 14, P1352–P1353 (2018).

52. Wolfsgruber, S., et al. Minor neuropsychological deficits in patients with subjective cognitive decline. Neurology 95, e1134–e1143 (2020).

53. Düzel, E., et al. CSF total tau levels are associated with hippocampal novelty irrespective of hippocampal volume. Alzheimer’s & dementia (Amsterdam, Netherlands) 10, 782–790 (2018).

54. Fischl, B., et al. Automatically parcellating the human cerebral cortex. Cereb Cortex 14, 11–22 (2004).

55. Iglesias, J.E., et al. A computational atlas of the hippocampal formation using ex vivo, ultra-high resolution MRI: Application to adaptive segmentation of in vivo MRI. Neuroimage 115, 117–137 (2015).

56. Desikan, R.S., et al. An automated labeling system for subdividing the human cerebral cortex on MRI scans into gyral based regions of interest. Neuroimage 31, 968–980 (2006).

57. Buckner, R.L., et al. A unified approach for morphometric and functional data analysis in young, old, and demented adults using automated atlas-based head size normalization: reliability and validation against manual measurement of total intracranial volume. Neuroimage 23, 724–738 (2004).

58. Ashburner, J. A fast diffeomorphic image registration algorithm. Neuroimage 38, 95–113 (2007).

59. Hessler, J., Jahn, T., Kurz, A. & Bickel, H. The MWT-B as an Estimator of Premorbid Intelligence in MCI and Dementia. Zeitschrift für Neuropsychologie 24, 129–137 (2013).

60. Washburn, R.A., McAuley, E., Katula, J., Mihalko, S.L. & Boileau, R.A. The physical activity scale for the elderly (PASE): evidence for validity. Journal of clinical epidemiology 52, 643–651 (1999).

61. Ganzeboom, H.B.G., De Graaf, P.M. & Treiman, D.J. A standard international socio-economic index of occupational status. Social Science Research 21, 1–56 (1992).

62. Wickham, H. ggplot2: Elegant Graphics for Data Analysis, (Springer-Verlag, New York, 2016).

63. Stuart, E., King, G., Imai, K. & Ho, D. MatchIT: Nonparametric Preprocessing for Parametric Causal Inference. Journal of Statistical Software 42 (2011).

64. Stuart, E.A. Matching Methods for Causal Inference: A Review and a Look Forward. Statistical Science 25, 1-21, 21 (2010).

65. Benjamini, Y. & Hochberg, Y. Controlling the False Discovery Rate: A Practical and Powerful Approach to Multiple Testing. Journal of the Royal Statistical Society. Series B (Methodological) 57, 289–300 (1995).

66. Aiken, L.S. & West, S.G. Multiple regression: testing and interpreting interactions., (Sage Publications, Newbury Park, CA, 1991).

67. Cohen, J., Cohen, P., West, S.G. & Aiken, L.S. Applied multiple regression/correlation analysis for the behavioral sciences, (L. Erlbaum Associates, Mahwah, NJ, 2003).

68. Hammers, A., et al. Three-dimensional maximum probability atlas of the human brain, with particular reference to the temporal lobe. Hum Brain Mapp 19, 224–247 (2003).

69. Brett, M., Anton, J.L., Valabrgue, R. & Poline, J.-B. Region of interest analysis using an SPM toolbox. Presented at the 8th International Conference on Functional Mapping of the Human Brain, June 2-6, 2002, Sendai, Japan. Neuroimage 13, 210–217 (2002).

