## Supplementary information for "Lifelong musical activity is associated with multi-domain cognitive and brain benefits in older adults"

### 1 SUPPLEMENT

#### 1.1 Supplementary methods

##### *1.1.1 Measurement of musical activity*

Musical activity across lifespan was assessed using the Lifetime of Experiences Questionnaire [1], as adapted for the German population (DZNE, Bonn) [2]. This self-reported questionnaire measures educational, occupational and cognitive lifestyle activities for three life periods (young adulthood: 13 – 30 years, mid-life: 30 – 65 years, and late-life: 65 years onwards). Within each life period, several complex specific and unspecific activities are assessed. Specific activities include those that are considered special to one particular life stage, such as education and occupation, while non-specific activities are measured for each life period and include frequency of participation in (1) playing a musical instrument, (2) social outings, (3) artistic activity (drawing, painting, writing), (4) physical activity, (5) reading, and (6) speaking a second language.

The frequency of musical activity was measured for each life period using the question (‘How often did you play an instrument?‘) with responses provided on a 6-point Likert scale (0 = ‘Never‘; 1 = ‘less than 1 time per month‘, 2 = ‘1 time per month‘, 3 = ‘2 times per month‘, 4 = ‘weekly‘, 5 = ‘daily‘). A recoding scheme was developed to reduce the information of the LEQ-item to the minimum that was needed to classify participants into the two groups: (1) a group with regular musical activity across the lifespan and (2) a control group that reported no musical activity during any life period. Three categories were used to recode participant responses for each life stage: For the category ‘no musical activity’, all answers ‘Never’ were coded as 0. For the category ’little musical activity’, answers ‘less than 1 time per month’ and ‘1 time per month’ were coded as 1. For the category ’frequent musical activity’, answers ‘2 times per month’, ‘weekly’ and ‘daily’ were coded as 2. Using these categories, participants were classified as having regularly engaged in musical activity across the lifespan, if they had a value of 2 for at least one life period, and no other lifespan was recoded as 0. Participants who received values of 0 for all life periods were classified as controls. Note, depending on a person’s age, classification was based on two (< 65 years of age) or three (≥ 65 years of age) life periods. To ensure that there were no false classifications, participants with missing responses on the musical activity item in any life period were excluded.

##### 1.1.2 Measurement of socio-economic status

The international socio-economic index of occupational information score ISEI, [3] was estimated using the occupational history of each participant, as assessed by the LEQ using 10 five-year intervals and following automated procedure. Occupational activities of each participant was used to assign O*Net codes (https://www.onetonline.org/) [4]. The O*Net codes were then transformed to Standard Occupational Classification codes (SOC), which were converted into codes of the International Standard Classification of Occupations (ISCO-08).The ISCO-08 codes were converted into ISEI scores using a crosswalk provided by Ganzeboom (retrieved from http://www.harryganzeboom.nl/ISCO08/index.htm, 04.2021). Finally, taking the total number of ISEI scores into account (maximum 10, according to the duration of occupational activity), an overall ISEI mean score was calculated per participant. This score was used as summary measure of socio-economic status.

#### 1.2 Supplementary results

##### Pre-analytical comparisons

Based on a prior population-based study [5], we expected significant differences between the two groups (lifelong regular musical activity and no musical activity) in demographic factors including known reserve proxies. In pre-analytical comparisons, the entire sample of participants with musical activity ($n$ = 73) and the control group were compared using baseline demographic, behavioral, neuropsychological, and neuroimaging variables. Independent Student's $t$-tests were used for all continuous variable and chi-squared ($\chi 2$) tests were applied for all categorical variables. In total, we identified 73 persons that reported lifelong musical activity and 356 participants with no musical activity (total: $n$ = 429) in the DELCODE baseline cohort. Our pre-analytical comparisons revealed significant group differences, with higher education, crystallized intelligence, SES, and both long-term and current physical activity found in those participants with a history of regular musical activity (all $p$'s < 0.05, supplementary Table 1). Groups were comparable in age, sex, and distribution of diagnostic groups (all $p$'s > 0.05). Finally, a matching procedure was applied based on relevant covariates, as derived in the pre-analytical comparisons, with the goal to identify a well-balanced control group [6,7].

=================

#### *Supplementary Table 1*

**Supplementary Table 1: Descriptive characteristics of the total neuroimaging DELCODE sample (*n* = 429)**

| | Musical activity | No musical activity | P value |
|---|---|---|---|
| Number (n) | 73 | 356 | - |
| Age (years) | 68.22 (6.60) | 69.44 (5.82) | 0.146 |
| Gender female/male (n) | 33/40 | 192/164 | 0.174 |
| Education (years) | 16.11 (2.75) | 14.28 (2.83) | < 0.001*** |
| Diagnostic group HC/FH/SCD (n) | 19/7/47 | 129/35/192 | 0.221 |
| SES [a] | 65.48 (16.82), *n* = 72 | 59.33 (17.64), *n* = 347 | 0.006** |
| Crystalized intelligence [b] | 33.29 (2.19), *n* = 72 | 32.08 (2.56), *n* = 353 | < 0.001*** |
| Physical activity, long-term [c] | 4.25 (0.77), *n* = 72 | 3.73 (1.13), *n* = 353 | < 0.001*** |
| Physical activity, current [d] | 33.94 (11.58), *n* = 69 | 30.60 (12.32), *n* = 346 | 0.033* |

Descriptive data are given if applicable as mean and standard deviation (in parenthesis). The actual sample size is provided, if different from sample size specified in first row. *P*-values correspond to independent *t*-tests for unequal variance with participant group as independent variable. Chi-square statistic was used to compare the distribution of categorical variables.
****p* < 0.001, ***p* < 0.01, **p* < 0.05.
***Key:*** HC, healthy control participants; FH, participants with family history of AD; GMV, gray matter volume; SCD, participants with subjective cognitive decline; SES, socioeconomic status.
[a] International socio-economic index (ISEI); [b] Multiple-Choice Vocabulary Intelligence Test (MWT); [c] Lifetime of Experiences Questionnaire (LEQ); [d] Physical Activity Scale for the Elderly (PASE).

==================

#### *Supplementary Figure 4*

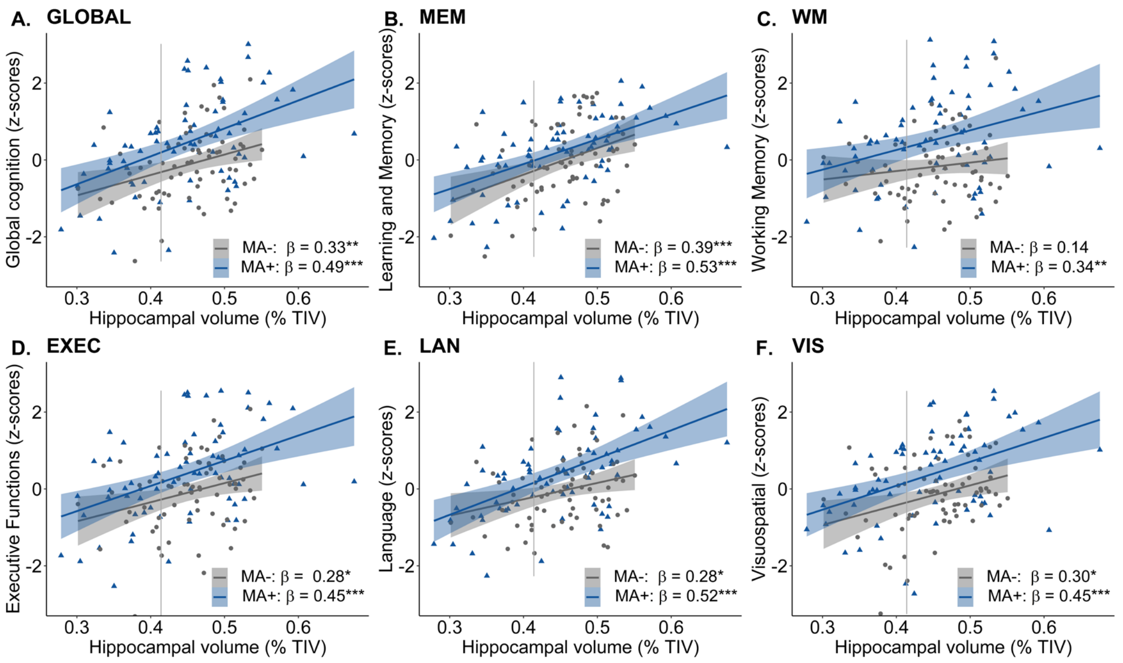


**Supplementary Figure 4: Moderation effect of lifelong musical activity in the hippocampal region.** There were no significant moderations effects of musical activity in the hippocampus. Global cognition (A) and domain-specific cognitive abilities (B-F) were positively associated with hippocampal volume for participants with lifelong musical activity (MA+, blue) and controls (MA-, gray). Scatter plots display the respective relationships. Individual data points, linear trends (solid lines), 95% confidence intervals (shaded areas), and standardized regression coefficients (β) within each group are provided. Gray vertical lines display the 90th percentile of the hippocampal GMV distribution in AD patients of the DELCODE study. Significance levels (uncorrected): ***$p < 0.001$, **$p < 0.01$, *$p < 0.05$. ***Key:*** GMV, gray matter volume; MA+, musical activity; MA-, no musical activity; TIV, total intracranial volume.

==================

#### *Supplementary Figure 6*

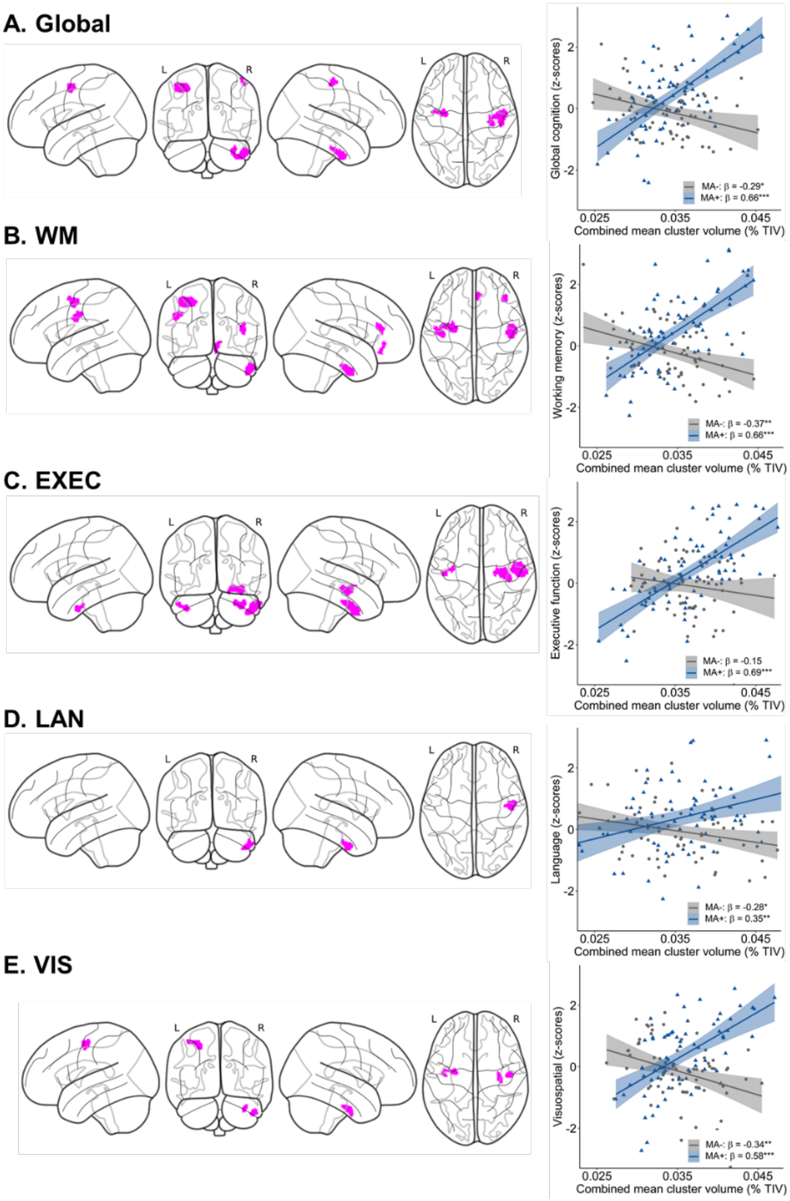


**Supplementary Figure 6: Associations between lifelong musical activity and regional volume distribution.** **A-E.** Results of the interaction analysis. The statistical maps display clusters ($p < .001$ uncorrected, color-coded in magenta) with a significant moderation effect of musical activity for global cognition (A, GLOBAL), working memory (B, WM), executive functions (C, EXEC), language (D, LAN), and visuospatial abilities (E, VIS). There was no significant interaction for the domain of learning and memory (data not shown). Statistical maps are depicted on a glass brain. Corresponding scatter plots show the associations using mean values extracted from the GMV maps in the combined cluster. Larger GMV in the combined cluster was associated with better cognitive abilities selectively in musically active participants (MA+, blue) compared to controls (MA-, gray). Individual data points, linear trends (solid lines), 95% confidence intervals (shaded areas), and standardized regression coefficients (β) within each musical activity group are provided. Significance levels (uncorrected): ***$p < 0.001$, **$p < 0.01$, *$p < 0.05$. ***Key:*** GMV, gray matter volume; TIV, total intracranial volume.

=================

#### 1.3 References


1. Valenzuela, M.J. & Sachdev, P. Assessment of complex mental activity across the lifespan: development of the Lifetime of Experiences Questionnaire (LEQ). *Psychol Med* **37**, 1015-1025 (2007).
2. Roeske, S.*, et al.* P3-591: A German version of the lifetime of experiences questionnaire (LEQ) to measure cognitive reserve: Validation results from the DELCODE study. *Alzheimer's & Dementia* **14**, P1352-P1353 (2018).
3. Ganzeboom, H.B.G., De Graaf, P.M. & Treiman, D.J. A standard international socio-economic index of occupational status. *Social Science Research* **21**, 1-56 (1992).
4. Peterson, N.*, et al.* Understanding Work Using the Occupational Information Network (O*NET): Implications for Practice and Research. *Personnel Psychology* **54**, 451-492 (2001).
5. Mansens, D., Deeg, D.J.H. & Comijs, H.C. The association between singing and/or playing a musical instrument and cognitive functions in older adults. *Aging & mental health* **22**, 964-971 (2018).
6. Zhang, Z. Propensity score method: a non-parametric technique to reduce model dependence. *Ann Transl Med* **5**, 7 (2017).
7. Ho, D., Imai, K., King, G. & Stuart, E. Matching as Nonparametric Preprocessing for Reducing Model Dependence in Parametric Causal Inference. *Political Analysis* **15**(2007).
